# Anti-Diffusion in an Algae-Bacteria Microcosm: Photosynthesis, Chemotaxis, and Expulsion

**DOI:** 10.1101/2023.12.14.571710

**Authors:** Praneet Prakash, Yasa Baig, François J. Peaudecerf, Raymond E. Goldstein

**Affiliations:** Department of Applied Mathematics and Theoretical Physics, Centre for Mathematical Sciences, University of Cambridge, Wilberforce Road, Cambridge CB3 0WA, United Kingdom; Institute de Physique de Rennes, Universite Rennes, UMR 6251, F-35000 Rennes, France

## Abstract

In Nature there are significant relationships known between microorganisms from two kingdoms of life, as in the supply of vitamin B_12_ by bacteria to algae. Such interactions motivate general investigations into the spatio-temporal dynamics of metabolite exchanges. Here we study by experiment and theory a model system: a coculture of the bacterium *B. subtilis*, an obligate aerobe that is chemotactic to oxygen, and a nonmotile mutant of the alga *C. reinhardtii*, which photosynthetically produces oxygen when illuminated. Strikingly, when a shaft of light illuminates a thin, initially uniform suspension of the two, the chemotactic influx of bacteria to the photosynthetically active region leads to expulsion of the algae from that area. This effect arises from algal transport due to spatially-varying collective behavior of bacteria, and is mathematically related to the “turbulent diamagnetism” associated with magnetic flux expulsion in stars.

---

In the early 1880s the biologist Theodor Engelmann performed experiments that were perhaps the first to use bacteria as sensors [1–3]. Several years prior he made the first observation of bacterial chemotaxis toward oxygen, by showing that putrefactive bacteria would migrate toward the chloroplasts of the filamentous alga *Spirogyra*. He then determined the “action spectrum” of photosyn-thesis—the wavelength-dependent rate of photosynthetic activity—by passing sunlight through a prism and projecting the spectrum onto a filamentous green algae held in a chamber that contained those self-same bacteria, which gathered around the algae in proportion to the local oxygen concentration, providing a direct readout of the oxygen production rate.

Although Engelmann’s system was engineered for a particular purpose, and at first glance involves a *one-way* exchange of oxygen for the benefit of bacteria, there are many examples of mutualistic exchanges between microorganisms from two distinct kingdoms of life. One of the most significant is that involving vitamin B_12_. In a landmark study [4], it was shown that a significant fraction of green algae that require this vitamin for their metabolism do not produce it, and as the ambient concentration of B_12_ in the aqueous environment is so low, they instead acquire it from a mutualistic relationship with bacteria, which benefit from a source of carbon.

The study of B_12_ transfer raises fascinating questions in biological physics related to the interplay of metabolite production, chemotaxis, and growth [5], including the issue of how organisms find each other and stay together in the turbulent environment of the ocean, and how advection by fluid flows arising from microorganism motility affects such mutualisms. As it is difficult to control the production of vitamin B_12_, we sought to construct a system in which the production of a chemical species needed by one member of an interacting pair of organisms could be controlled by the experimentalist. Taking motivation both from Engelmann’s experiments and the B_12_ system, we introduce here a coculture [6] in which an obligate aerobic bacterium (one that requires oxygen) that is chemotactic toward oxygen, coexists with a green alga whose photosynthetic activity can be turned on and off simply by controlling the external illumination.

We use the bacterium *Bacillus subtilis*, whose aerotaxis has been central in the study of bioconvection [7] and in the discovery of “bacterial turbulence” [8], the dynamical state of a concentrated suspension with transient, recurring vortices and jets of collective swimming on scales large compared to the individual bacteria. The alga species is the well-studied unicellular *Chlamydomonas reinhardtii*, a model organism for biological fluid dynamics [9] with readily available motility mutants useful in probing the role of swimming in metabolite transfer. Together these define what we term the *algae-bacteria-chemoattractant system* (ABC). While coupled population dynamics problems have been studied from the familiar reaction-diffusion point of view in bacterial range expansion [10], marine [11] and more general ecological contexts [12], we show that there are physical effects that go beyond that standard level of treatment. Chief among them is the way in which collective motion can act as a “thermal bath” in enhancing the diffusivity of suspended particles [13, 14], which in turn raises fundamental issues concerning generalizations of Fick’s law [15, 16].

A disarmingly simple experiment to understand the dynamics of the ABC system involves a thin, quasi-two-dimensional suspension of non-motile algae and fluorescently labelled bacteria at initially uniform concentrations. As depicted in Fig. 1, a shaft of photosynthetically active light is cast on the suspension, triggering oxygen production by the illuminated algae. As shown in Figs. 2(a,b), this leads to chemotaxis of bacteria into the illuminated region, producing a high concentration of bacteria. Remarkably, we find that algae are then expelled from the illuminated region. Quantitative measurements of the local bacterial dynamics in the the system show that this expulsion is associated with a *gradient of collective bacterial behavior* from its peak at the center, leading to a outward algal transport. On longer time scales, after the algal expulsion, the bacterial concentration returns to uniformity as the bacteria diffusive outward in the absence of chemotactic stimulus. We name this *Type I* dynamics. Figures 2(c,d) show that at sufficiently high initial bacterial concentrations a new, *Type II* behavior is observed; expulsion of alga and consumption of oxygen are sufficiently rapid that many bacteria in the illuminated region become hypoxic, transition to an immotile state, and are then also expelled into the dark.

**FIG. 1.**
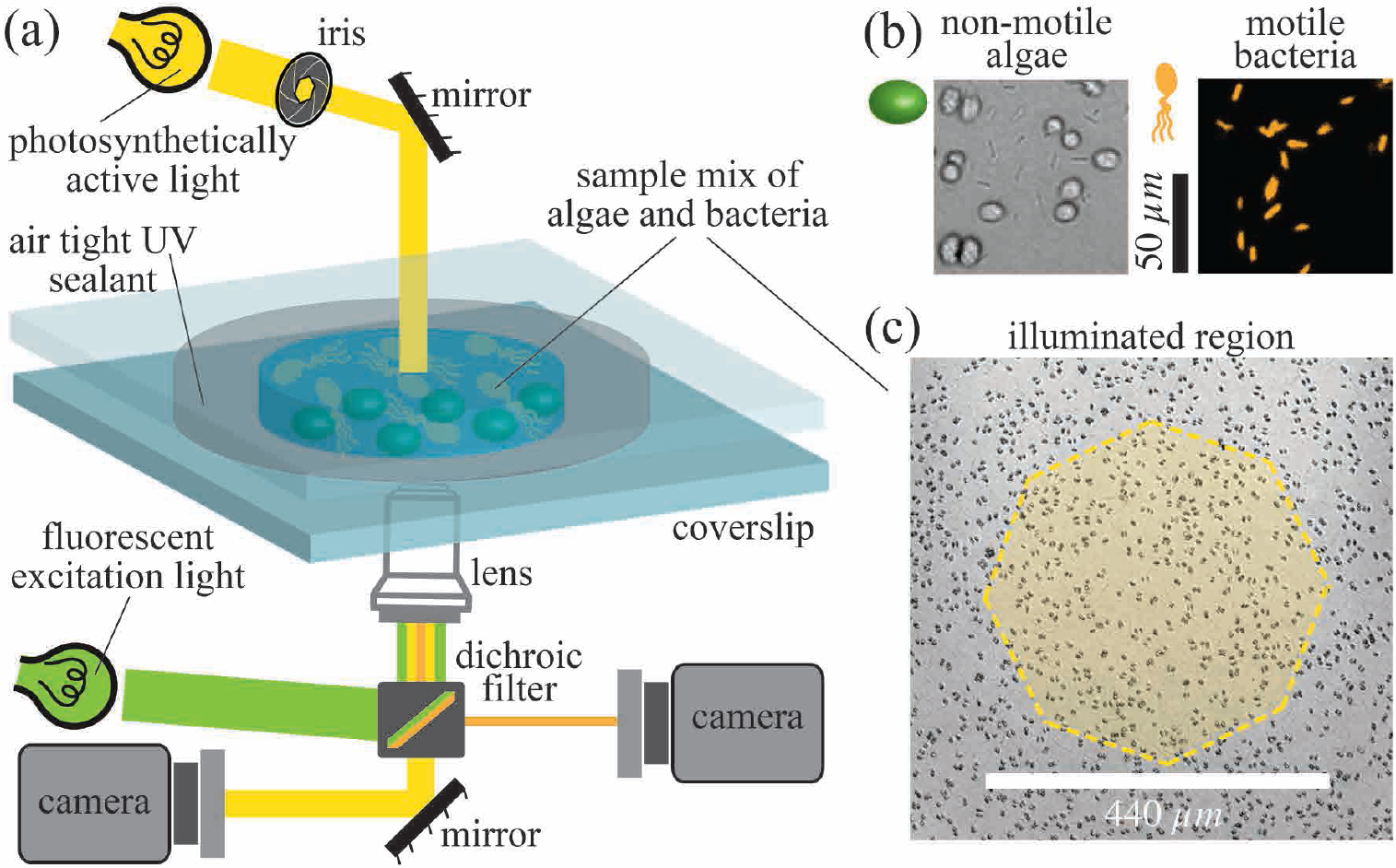
Experimental setup. (a) Schematic of system to illuminate a coculture of a fluorescent strain of the bacterium *B. subtilis* and a flagella-less mutant of the green alga *C. reinhardtii*, initially distributed uniformly within the sample chamber. After a shaft of visible light illuminates the central region the algae produce oxygen to which the bacteria are attracted. (b) Algae and bacteria as observed through the brightfield and fluorescent channels, respectively. (c) Yellow shading indicates the illuminated region, with dark algae visible in the background.

**FIG. 2.**
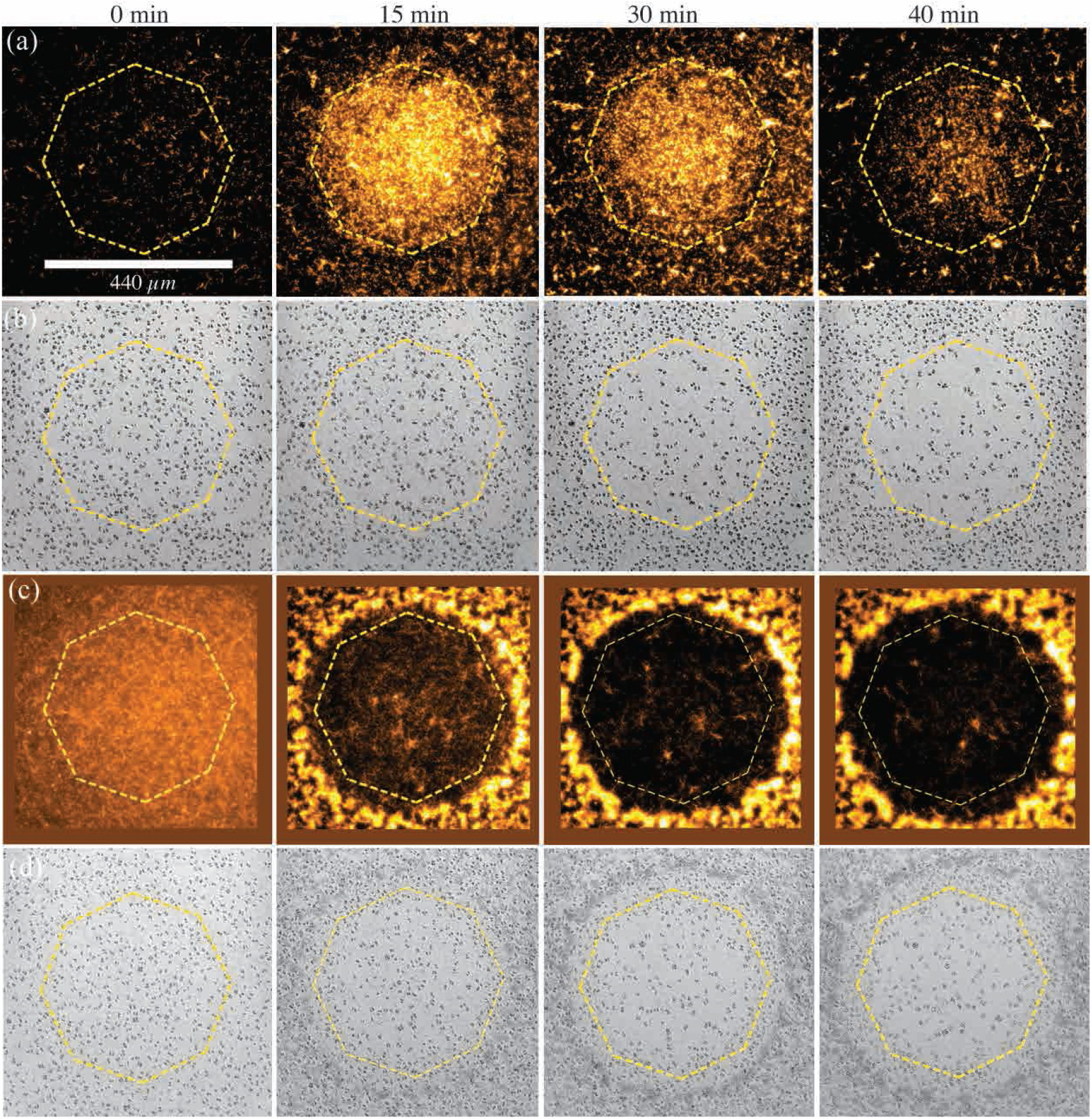
Bacterial influx and algal expulsion. (a) Type-I dynamics. Spatio-temporal evolution of bacterial concentration after illumination is initiated, where octagonal ring indicates boundary of illuminated region. At the low initial bacteria concentration (*∼* 1 *×* 10^8^ cm^*−*3^), bacteria first move into the illuminated region and then retreat. (b) Algal concentration as a function of time, demonstrating expulsion from illuminated region. (c) Type II dynamics. At a higher concentration (*∼* 5 *×* 10^8^ cm^*−*3^), many bacteria become non-motile and are expelled into the dark region, forming a concentrated circular accumulation that acts as a natural boundary. (d) Algae are also expelled, accumulating inside the confines of the bacterial accumulation.

We develop here a system of coupled PDEs that describes the ABC system that provides a quantitative account of the experimental observations. These PDEs incorporate diffusion, chemotaxis, oxygen production and consumption and algal transport by collective effects. The expulsion of algae is similar to well-known processes in magnetohydrodynamics termed “flux expulsion”. First discovered for magnetic fields in a prescribed constant vortical flow field [17], in which field lines are expelled from the vortex, it was later recognized that this expulsion can arise from gradients in the intensity of turbulence whose random advection of the magnetic vector potential leads to a gradient in its effective diffusivity and thence to “turbulent diamagnetism” [18].

## Methods

The coculture used strain 168 of *B. sub-tilis* which was genetically engineered to express yellow fluorescent protein m-Venus with excitation at 515 nm and emission at 528 nm [19]. A single bacterial colony was picked from an agar plate and grown overnight in Terrific Broth (TB) on an orbital shaker at 240 rpm and 30 ^*°*^C. The bacteria intended for experiments were grown from overnight culture until exponential growth phase in Tris-min medium spiked with TB medium (Tris-min + 0.1 % w/v glycerol + 5 % w/v TB). The non-motile *C. reinhardtii* strain CC477 (*bld1-1*) was sourced from the Chlamydomonas Resource Center [20] and grown in Tris-min medium, on an orbital shaker at 240 rpm and 20 ^*°*^C. The diurnal cycle was 12 h cool white light (*∼* 15 *µ*mol photons/m^2^s PAR), and 12 h in the dark.

Prior to experiments, bacteria and alga from their respective exponential growth phases were mixed in a modified Tris-min medium, with glycerol as a carbon source and bovine serum albumin to prevent cell adhesion (Tris-min + 0.1 % w/v glycerol + 0.01 % v/v BSA). The desired number density of cells was achieved by centrifugation at 4000 g. All cell concentrations and sizes were measured using a Beckman Coulter Counter (Multisizer 4e). The concentration of algae was fixed at 5 *×* 10^6^ cm^*−*3^ while varying the bacteria concentration in the range (0.5 *−* 5) *×* 10^8^ cm^*−*3^. The mixture of cells for each experiment was transferred to a glass cover slip chamber with a depth of 300 *µ*m, separated by double-sided tape, and sealed airtight using UV glue. The chamber surfaces were passivated with PEG (*M*_*w*_ = 5000 g/mol).

Experiments were performed on a Nikon TE2000-U inverted microscope. The spatio-temporal variation in bacterial concentration was monitored by epifluorescence illumination with a *×*20 objective using a highly sensitive, back-illuminated camera (Teledyne Prime Σ 95B). Movies of algae cells were recorded through the brightfield channel using a Phantom V311 high-speed camera (Vision Research) at *×*20 magnification. The halogen lamp served as both a brightfield and photosynthetic light source. The size of a light shaft used to trigger photosynthesis was controlled by the field iris in the microscope condenser arm, producing an octogonal boundary in the focal plane with mean radius of *R* = 220 *µ*m.

### Experimental Results

Figures 2 and 3 summarize the main experimental observations associated with a homogeneous initial condition in darkness that is then illuminated with a shaft of light. For an initial bacterial concentration of *b* = 1 *×* 10^8^ cm^*−*3^ giving Type I dynamics, we observe over the course of the first *∼* 10 min after the start of illumination that the concentration of bacteria at the center of the illuminated region increases dramatically, as visualized in Fig. 1(a) and quantified in Fig. 2(b), reaching a peak enhancement of a factor of *∼* 5, with a roughly linear decrease out to the edge. That peak then relaxes away until the bacterial concentration is again nearly uniform after *∼* 35 min. At the peak of accumulation, after *∼* 15 min, there is a clear bacterial depletion zone just outside the illuminated region, whose width is estimated to be *∼* 150 *µ*m. During this period, as shown in Fig. 2(b), the algal concentration becomes strongly depleted in the illuminated region, leading to a ring of accumulation at the boundary.

**FIG. 3.**
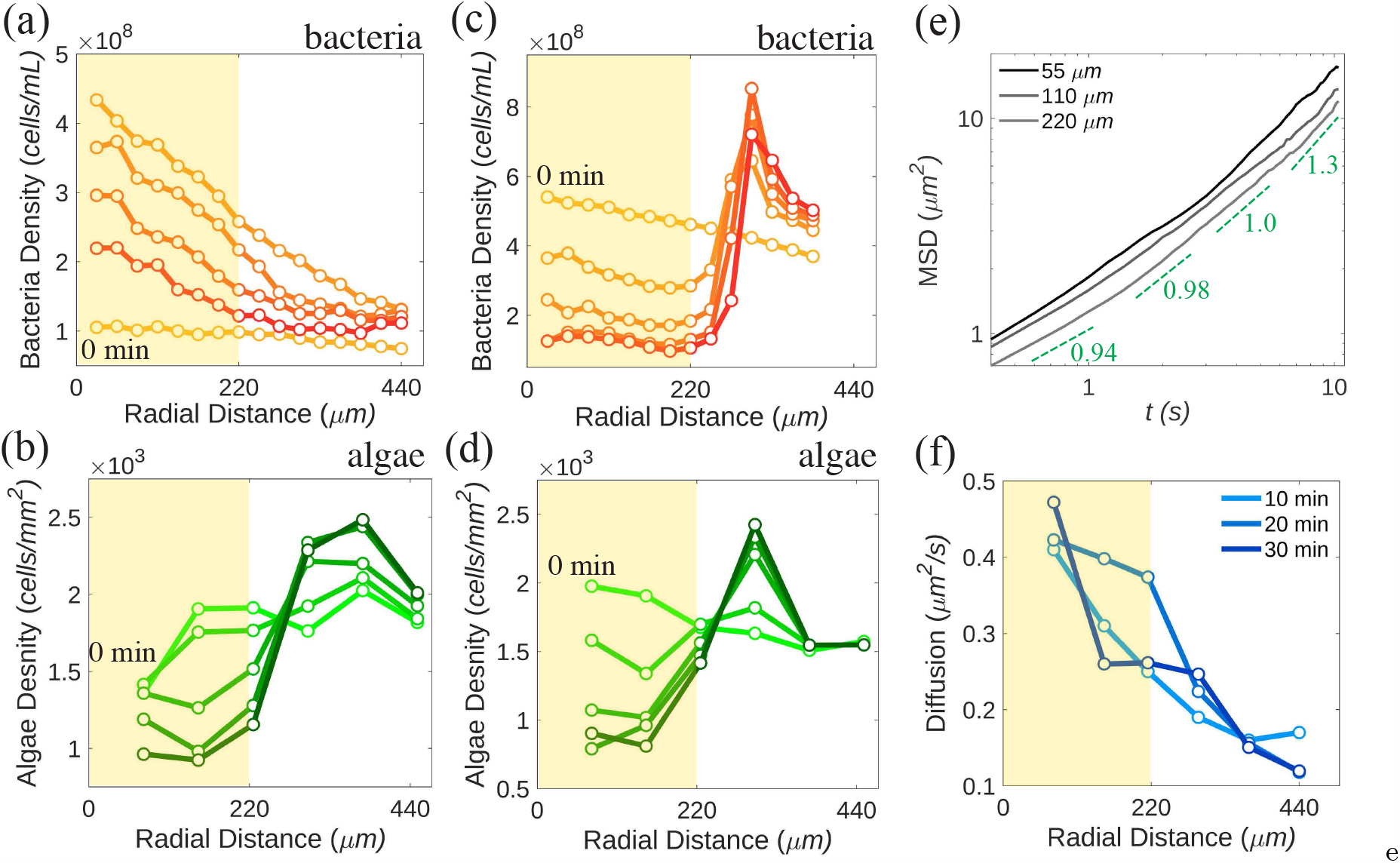
Bacterial accumulation and algal expulsion. Yellow shading in the plots up to 220 *µ*m indicates illuminated region. (a) Bacterial and (b) algal concentrations for a coculture with initial concentrations *b* = 1 *×* 10^8^ cm^*−*3^ and *a* = 5 *×* 10^8^ cm^*−*3^, showing Type I dynamics. Shading of symbols and lines increase with time, with a ten minute interval between data sets. (c,d) As in (a,b) but with initial bacterial concentration *b* = 5 *×* 10^8^ cm^*−*3^, exhibiting Type II dynamics. (e) MSD of algae versus time for the case (a,b) at different radii in the illuminated region. Dashed lines indicate apparent slope at different times. (f) Algal diffusivity determined in linear regime of MSD versus radial distance at various times after start of illumination.

We measured the mean squared displacement (MSD) of algae versus time at a range of radii *r* from the light shaft center. The MSD in Fig. 3(e) exhibits systematic upward curvature, with a local exponent of unity at time *t*_1_ *∼* 6 *−* 7 s, but faster behavior for *t > t*_1_. Such superdiffusion for tracers is a well-known consequence of active turbulence in concentrated bacterial suspensions [8, 13]. A heuristic illustration of the strong gradient in collective behavior in the illuminated region is obtained by determining an effective algal diffusion constant 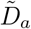 from the slope of the MSD curve at *t*_1_ (Fig. 2(f)), which is a strongly decreasing function of distance from the shaft center, that mirrors the decreasing bacterial concentration. The largest of these diffusivities is *∼* 25 times the purely thermal value *D*_th_ = *k*_*B*_*T/*6*πµa ≃* 0.02 *µ*m^2^/s, where *µ* is the medium viscosity and *a* = 10 *µ*m is twice the algal radius since most algae exist as pairs (“palmel-loids”). Yet, by itself, an effective algal diffusivity 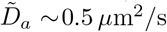 implies a diffusive time 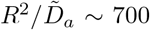 min for algae at the center to escape purely by random motion, far longer than the observed time of 30 min; the superdiffusive behavior seen for *t > t*_1_ in Fig. 3(e) signals a qualitatively different transport process.

At the higher initial bacterial concentration *b* = 5 *×* 10^8^ cm^*−*3^ we observe the distinct Type II behavior: expulsion happens much more rapidly, the bacterial concentration profile becomes nonmonotic in radius, with a strong peak just outside the illuminated region, and the peak of the concentration of expelled algae is narrower. Close microscopic inspection of the region of high bacterial concentration in the dark region shows that most of the bacteria there are immotile and the expelled algae reside just inside the bacterial accumulation ring. We deduce that many bacteria in the illuminated region have become hypoxic and are expelled from that area through much the same process that expelled the algae.

### The ABC model

We now turn to a mathematical model for the behavior described above, focusing first on Type I expulsion. In this case, a minimal description of the system entails the concentration fields *a*(**r**, *t*), *b*(**r**, *t*) and *c*(**r**, *t*) for algae, bacteria, and oxygen, respectively. If *D*_*c*_ is the oxygen diffusion constant (assumed independent of all other variables), *k*_+_ is the rate of oxygen production per algal cell when illuminated with a spatially varying light intensity *I*(*r*) and the oxygen consumption rate per bacterium has a Michaelis-Menten form, then

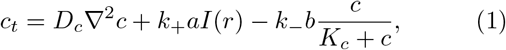

where *K*_*c*_ is the Michaelis constant.

Consider first the situation of uniform illumination (*I* = 1) and uniform concentrations *a*_0_ and *b*_0_ of algae and bacteria. A steady state *c*^*\**^ = *K*_*c*_*k/*(1 *− k*), with *k* = *k*_+_*a*_0_*/k*_*−*_*b*_0_, can be reached in which consumption balances production, provided *k <* 1. We assume this inequality is always satisfied and typically invoke the *weak production limit k ≪* 1, in which the oxygen consumption can be approximated as *k*_*−*_*bc/K*_*c*_.

We explain algal expulsion from the illuminated region through three intermediate calculations; oxygen production in a uniform suspension; bacterial chemotaxis in the presence of oxygen production; algal dynamics due to an inhomogeneous bacterial concentration. Suppose that the light is constrained to a shaft of radius *R*. Scaling time, space, and concentrations via *T* = *tD*_*c*_*/R*^2^, *η* = *r/R, χ* = *c/c*^*\**^, *α* = *a/a*_0_, and *β* = *b/b*_0_, letting *ϵ* = *k/*(1 *− k*), and introducing the *screening length*

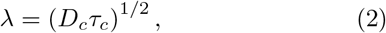

where *τ*_*c*_ = *K*_*c*_*/k*_*−*_*b*_0_ is the characteristic consumption time of oxygen, the dynamics (1) takes the form

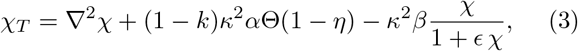

where now 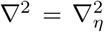, and Θ is the Heaviside function. Here, *κ*^2^ = *R*^2^*/λ*^2^ is also the ratio *τ*_*D*_*/τ*_*c*_ of the time *τ*_*D*_ = *R*^2^*/D*_*c*_ *∼* 25 s for oxygen to equilibrate diffusively across the illuminated region to the consumption time.

If we clamp concentrations *α* and *β* at unity, take *k ≪* 1, and enforce continuity in *χ* and *χ*_*η*_ at *η* = 1, the steady state of (3) in an unbounded domain is

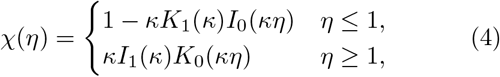

in terms of modified Bessel functions *K*_*ν*_ and *I*_*ν*_. From the inner solution we see that the oxygen concentration at the centre of the illuminated region asymptotes to unity (*c*^*\**^ in unrescaled units) for large domain size *κ*, but is attenuated strongly as *κ* falls below unity. The outer solution behaves as *χ ∼* exp(*−*(*r − R*)*/λ*), showing that *λ* serves as the characteristic penetration depth of oxygen into the surrounding bacterial population and sets the depletion zone seen in Fig. 1(b). With *D*_*c*_ *∼* 2 *×* 10^3^ *µ*m^2^/s and *R* = 220 *µ*m, and the estimate *τ*_*c*_ *∼* 10^2^ s [21], we find *λ ∼* 400 *µ*m, and thus *κ ∼* 1 *−* 2.

Relaxing the assumption of uniform bacterial concentration, it is clear that beyond the time *τ*_*D*_ bacteria within a distance *λ* of the edge of the illuminated region will experience the steepest oxygen gradient and chemotax most rapidly inwards. This can be described by the simplest combination of diffusion and chemotaxis as in the Keller-Segel model, [22], *b*_*t*_ = *D*_*b*_*∇*^2^*b −* ***∇*** · (*gb****∇****c*) where the diffusion constant *D*_*b*_ arises from random cel-lular swimming and the response coefficient *g* is taken to be constant. Scaling as above, we obtain

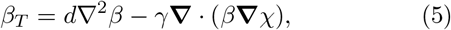

where *d* = *D*_*b*_*/D*_*c*_, and *γ* = *gc*^*\**^*/D*_*c*_.

A steady state can be reached when the chemotactic flux *γβ****∇****χ* balances the diffusive flux *d****∇****β*, yielding

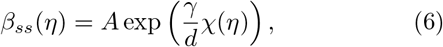

where *A* is a normalization constant. While this steady state profile captures the observation that the bacterial concentration reaches its maximum at *η* = 0, where *χ* itself is maximized, the steady state profile (6) only develops on time scales sufficient for bacteria outside the depletion zone to move inwards and replenish partially the depletion. This time will be at least (*R* + *λ*)^2^*/D*_*b*_ ≫ *τ*_*D*_. Prior to this the bacterial concentration is nonmonotonic, with the depletion zone seen in Fig. 2(a).

The flagella-less algae used in our experiments do not swim; their movement arises from collective flows driven by the concentrated bacteria. At the very least this leads to enhanced diffusivity, which, as seen in Fig. 3(f), varies in space, and by itself, is captured by an algal flux 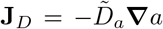. The appearance of “bacterial turbulence” at higher concentrations implies that the algae are passive scalars in an *inhomogeneous* turbulent flow. It is well-known that passive particles will be expelled into separatrices between regions of high vorticity. When those boundaries change with time, the effect to transport particles from regions of high turbulence to low, without requiring any gradients in particle concentration, akin to *chemokinesis*, where cells accumulate in regions where they swim slowly. In a simple approximation, the extent of the turbulent transport is proportional to the bacterial concentration, giving a contribution to the algal flux **J** = *−pa****∇****b* for some *p >* 0. With the concentration seen in Fig. 3(a), whose radial gradient is negative, this contribution leads to an *outward* flux of algae.

Assembling these contributions and rescaling as above, we obtain two equivalent forms of the dynamics,

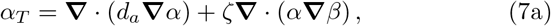

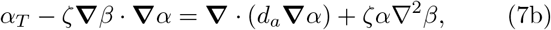

where 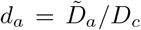 and *ζ* = *pb*_0_*/D*_*c*_. In the first form (7a), we see a parallel to the bacterial chemotaxis equation (5); algae exhibit negative “bacteria-taxis”. In symmetrizing the two problems, this “completion” of the dynamics is loosely analogous to the introduction of the displacement current in Maxwell’s equations as a parallel to Ampere’s law. In the second form (7b), we see that there is an explicit advective contribution and a “reactive” term *α∇*^2^*β*. This latter contribution plays an important role in the algal expulsion, as the bacterial concentration within the illuminated region has a large negative second derivative, and this term thus forces *α* down there, whereupon it is advected outward. This is a process of “anti-diffusion”.

Figures 4(a,b) show how the model defined by Eqs. (3), (5) and (7b) provide a quantitative fit to the observed Type I dynamics. In particular, we see the prompt accumulation of bacteria in the illuminated region followed by expulsion of algae. In the model and in experiment, the fraction of algae ultimately expelled is *∼* 0.5, so there is still oxygen production within the illuminated region after *∼* 30 min, leading to continued chemotactic attraction of bacteria inward. This leads to a slower decay of the bacterial concentration back to its original uniform, low value than seen in experiment. This may reflect processes such as bacterial adaptation to the oxygen and a gradual reduction in oxygen production by algae.

**FIG. 4.**
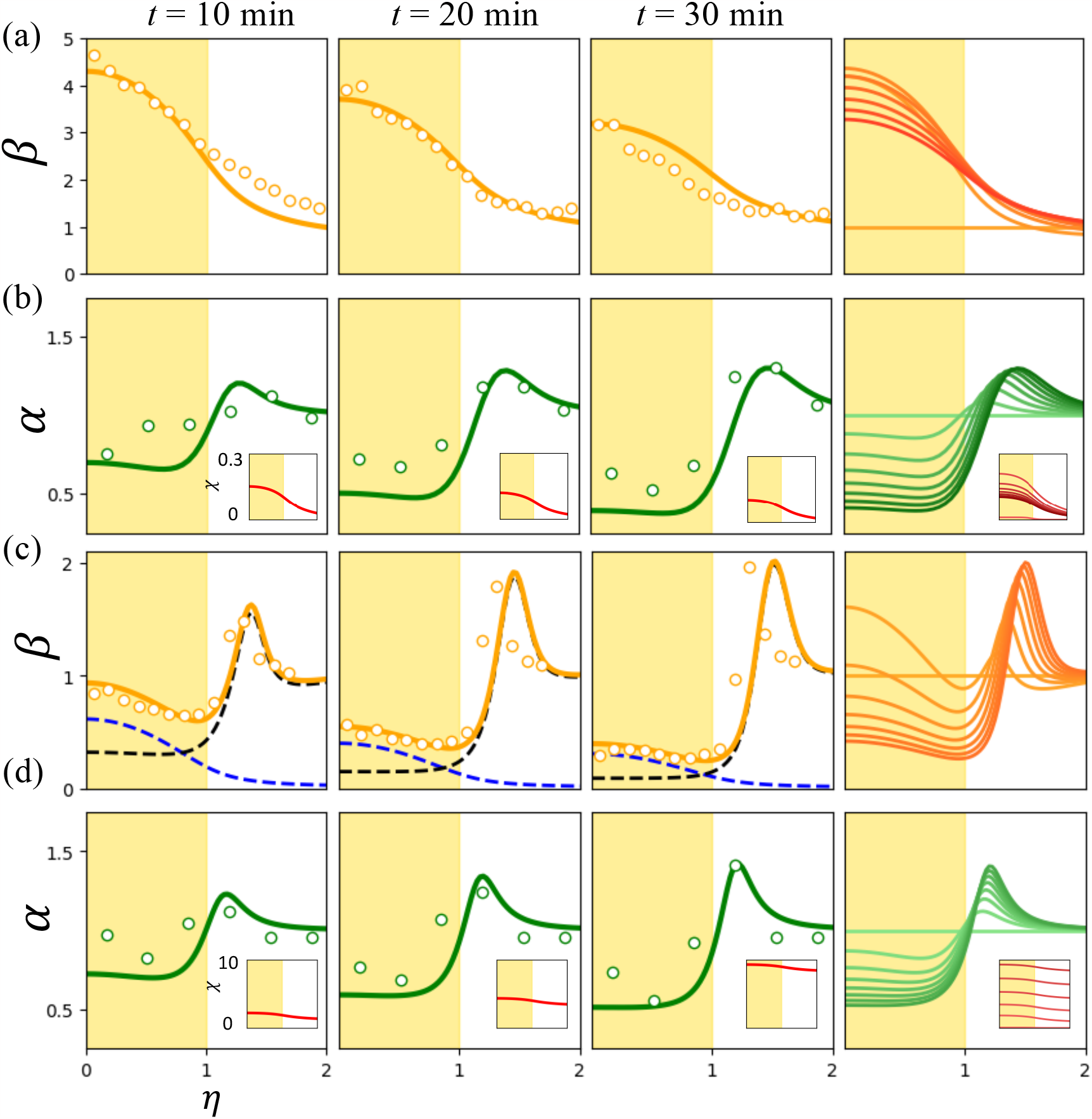
Theoretical predictions. (a,b) Type I dynamics. (a) Non-dimensional bacterial dynamics from experiment (circles) compared to theoretical fit (solid lines) for first thirty minutes of experiment. Rightmost column in each row shows the theoretical time evolution over 30 minutes, sampled every *∼* 3 minute. Shading intensity increases with time. (b) As in (a) but for algae, showing expulsion. Insets show theoretical oxygen profiles *χ*. (c,d) Type II dynamics. (c) Experimental bacterial dynamics (circles) compared to predictions of the ABCD model. Dashed lines indicate immotile (black) and active (blue) bacteria, whose sum is orange solid line. Rightmost plot shows time evolution of the total bacterial concentration. (d) Algal expulsion for high density experiment. For (a),(b) we took *κ*^2^ = 1, *ϵ* = 1, *d*_*b*_ = 0.1, *d*_*a*_ = 0.0001, *γ* = 1.2, *ζ* = 0.002. For (c), (d), we use *γ* = 0.35, *ζ* = 0.005, *κ*^2^ = 5, *d*_*δ*_ = 10^*−*4^, *ρ* = 1.0, *χ*^*\**^ = 0.1, *ζ*_*d*_ = 0.04, *δ*^*\**^ = 0.01 with all other parameter values held over.

**FIG. 5.**
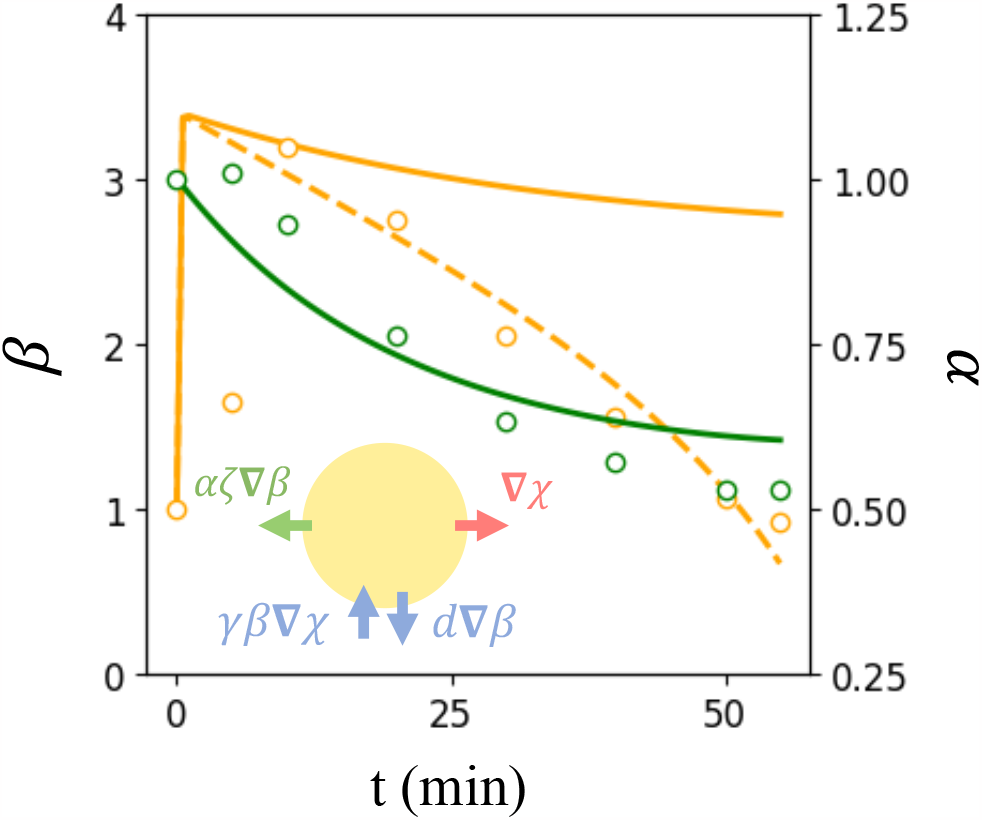
Spatially averaged dynamics. Average concentration of bacteria (orange) and algae (green) in illuminated region from experiments (circles) and model (lines) shown in scattered points. For bacteria, simple chemotaxis is shown by solid orange line, adaptive chemotaxis shown dashed. Inset: Schematic of fluxes across the illuminated region.

### The ABCD model

While the ABC model can account for the essential features of Type I dynamics, it does not allow for loss of motility of bacteria at low oxygen concentrations in Type II dynamics. This transformation has been recognized as important in the context of bioconvection [21, 23], where the influx of oxygen at the air-water interface of a bacterial suspension competes with consumption within the fluid, leading to a hypoxic region hundreds of microns below the surface. Hypoxia-induced motility transitions have also been observed in the penetration of oxygen into suspensions of *E. coli* [24].

To account for this transformation we view the “dormant”, non-motile state of the bacteria, with concentration *d*, as a separate population distinct from the motile form, so the extended “ABCD” model involves algae, bacteria, chemoattractant, and dormant bacteria. The interconversion rate as a function of oxygen concentration *c* is taken as a simple generalization of the substrate-dependent growth of the Monod model, *v*_*con*_*bK*_*sat*_*/*(*c* + *K*_*sat*_), where *v*_*con*_ is the maximum conversion rate to the immotile form and *K*_*sat*_ is the concentration at which half-maximal conversion occurs.

The dynamics of dormant bacteria include generation, expulsion, and a very small diffusion constant *D*_*d*_. Setting *δ* = *d/b*_0_, the rescaled dynamics takes the form

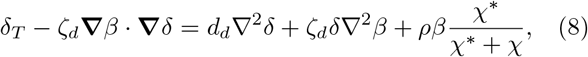

where *d*_*d*_ = *D*_*d*_*/D*_*c*_, *ρ* = *v*_*con*_*R*^2^*/D*_*c*_ and *χ*^*\**^ = *K*_*sat*_*/c*^*\**^. Accordingly the bacterial dynamics (5) acquires the corresponding loss term from conversion, becoming

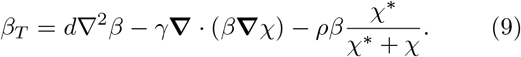

A final modification to the ABC model involves steric effects on algal diffusion that occur when the concentration of dormant bacteria is large. This is modelled by modifying the rescaled algal diffusivity *d*_*a*_ to *d*_*a*_(1 *− δ/*(*δ* + *δ*^*\**^)), a form that reduces *d*_*a*_ at large *δ* but retains its positivity. With these components, Fig. 4(c,d) show that the ABCD model captures the expulsion of both algae and immotile bacteria, with a narrow accumulation of dormant bacteria blocking transport of algae.

### Spatially-averaged model

Insight into the basic process of algal expulsion can be obtained by constructing a spatially-averaged model which takes as dynamical degrees of freedom the mean concentrations of algae, bacteria, and oxygen inside the illuminated region, denoted by 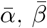, and 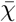. As indicated schematically in Fig. 5(a), the specification of this averaged model requires estimates of the various fluxes across the boundary of the illuminated region. We anticipate that turbulent bacterial dynamics will homogenize oxygen inside the illuminated region, save for the area close to the boundary of width *λ* where the concentration rapidly decreases. From an integral form of (1) and the divergence theorem we have 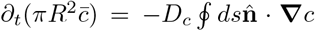, where 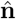 is the out-ward normal to the illuminated region. If we estimate 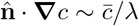, then the PDE (1) becomes the ODE

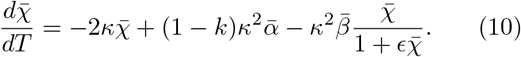

Similar estimates hold for the averaged bacteria dynamics, where the relevant fluxes are the outward diffusive and inward chemotactic contributions. The diffusive contribution is precisely analogous to that for oxygen above. Since on the time scales of the experiment only bacteria within a distance *λ* outside the illuminated region sense the gradient, the inward Keller-Segel flux is also estimated as above, but with an upper limit on the bacterial concentration due to steric effects. This leads to the ODE version of (5),

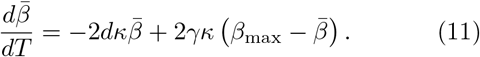

Finally, the algae have an active flux *−pa****∇****b* that leads to expulsion, but there is little back-diffusion into the illuminated region. Yet, as algae accumulate at the boundary (Fig. 2), they form a thick ring that inhibits continued expulsion of algae. Introducing a saturation of the algal flux we obtain the ODE version of (7a),

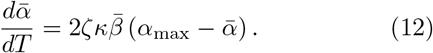

Using parameter values taken from Fig. 4 and fitting *β*_max_ ≈ 5 and *α*_max_ ≈ 0.5, we obtain the solid green and orange curves shown in Fig. 5 for mean algal and bacterial concentrations within the illuminated region. The reduced model captures the slow exponential decay of the algal concentration, but while it reproduces the rapid bacterial influx in the first few minutes of the experiments, the predicted decay of the bacterial concentration, as remarked earlier for the full ABC model, is far slower than observations. The possibility that this discrepancy arises from adaptation of the bacteria to elevated oxygen levels can be explored by assuming a simple linear decay of the chemotactic coefficient, as *γ*(*T*) = *γ*_0_ *−T/τ*, where *τ* sets the adaptation rate. For *τ* = 1200, the bacterial dynamics (dashed orange line in Fig. 5) matches closely with data, while the algal expulsion (not depicted) is essentially unchanged from the non-adaptive case.

## Discussion

As our results suggest new phenomenology in active matter systems, it is instructive to return to the connection between algal expulsion and flux expulsion in magnetohydrodynamics. In the original case considered [17], a magnetic field in a fluid with velocity **u** obeys the Maxwell equation,

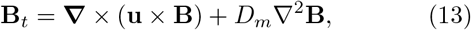

where *D*_*m*_ is the magnetic diffusivity, and both **B** and **u** are taken to confined to the *xy*-plane. The magnitude of the magnetic vector potential 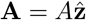 then obeys the advection-diffusion equation

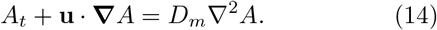

With **u** a prescribed single-vortex velocity field, the initial condition *A* = *B*_0_*x*, corresponding to the space-filling uniform magnetic field **B** = ***∇*** *×* **A** = *−B*_0_**ŷ**, homogenizes inside the vortex, leaving **B** = 0 zero there but *A* essentially undisturbed outside.

In the case of turbulent transport consider later [18], it was shown that on long time and length scales the effect of the inhomogeneous turbulence is captured by an equation of motion for a vector potential component 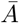, averaged on those scales, of the form

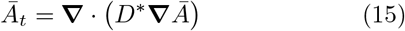

where *D\** is a turbulent diffusivity tensor. For the particular case in which the turbulence varies along one direction, say *y*, then the averaged *x*-component of the magnetic field 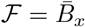 obeys

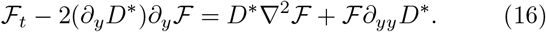

This form has the same structure as our Eq. (7a), with the bacterial concentration *β* playing the role of the diffusivity tensor component. We conclude that the main difference between the two problems is the expulsion of a scalar (concentration) field in the present context compared to the expulsion of a vector quantity in MHD.

The results described here highlight the rich dynamical behavior that occurs in mixed active matter systems involving microorganisms from two Kingdoms of Life. While the dynamics of symbiosis is generally studied on time scales relevant to population dynamics or evolution [25–27], our findings indicate that short-term dynamics on the scale of minutes and hours can have a considerable impact on the spatial-temporal aspects of association, where chemotaxis and phenotypic switching dominate. Although we considered the simple case of light intensity that is piecewise constant, inhomogeneous activity arises. The treatment of such nonuniform active matter has received attention only recently, in the context of “invasion” [28] and active gels [29], but is central to any discussion of realistic ecologies. In this sense, generalizing the present setup to allow algal motility and phototaxis in the presence of an inhomogeneous light field may reveal even more striking dynamics when the oxygen sources are themselves motile.

We are grateful to Jim Haseloff for providing the fluorescent strain of *B. subtilis* and to Kyriacos Leptos for sumerous discussions. This work was supported in part by the Gordon and Betty Moore Foundation, Grant No. 7523 (PP & REG) and the U.K. Marshall Aid Commemoration Commission (YB).

